# Meat the Fat: From High Lipid Content to Nutritional Enhancement to Reconstruct Wagyu Fat with Tunable Rheology and Organoleptic Fidelity for Cultivated Meat

**DOI:** 10.64898/2025.12.08.692892

**Authors:** Fiona Louis, Ais Cizeron, Mai Furuhashi, Shoji Takeuchi, Tatsuya Nojima, Shiro Kitano, Michiya Matsusaki

**Author notes:** Corresponding authors: Fiona Louis Michiya Matsusaki.

## Abstract

Cultivated meat holds promise as a sustainable and ethical alternative to conventional meat, but reproducing the rich flavor, texture, and lipid composition of premium beef, such as Wagyu, remains a major hurdle. In this study, we redefine bovine adipogenesis by systematically optimizing edible biomaterials, fatty acid delivery systems, and culture conditions to engineer high-fat cultivated tissues with authentic sensory properties. Among various candidates, 1% alginate emerged as the optimal scaffold, providing mechanical softness compatible with adipogenic differentiation and mimicking the rheological properties of native fat. Supplementation with oleic acid complexed to edible albumins, especially whey protein (WP), significantly enhanced lipid accumulation and monounsaturated fatty acid content, particularly oleic acid, which reached up to 80.6% of total fatty acids. Engineered fat tissues displayed soft, spreadable textures and organoleptic characteristics consistent with high-quality beef fat. Fatty acid profiling revealed a bovine-like omega-6 signature, dominated by linoleic, arachidonic, and dihomo-γ-linolenic acids, alongside detectable omega-3 subtypes including alpha-linolenic acid and trace levels of EPA and DHA. This composition not only validates the physiological relevance of the engineered fat, but also highlights its potential to deliver bioactive lipids rarely found in terrestrial animal products. Additionally, the WP-oleic acid system supported robust adipogenesis even under lower serum conditions. These findings establish a food-grade, tunable platform for producing physiologically and nutritionally enhanced fat tissues, advancing the realism, functionality, and health potential of next-generation cultivated meat.

## Introduction

Recent advancements in the field of cultivated meat have highlighted the potential for laboratory-based meat production to address pressing global challenges, including environmental sustainability and ethical considerations associated with traditional livestock farming [1–3]. While substantial progress has been made in reconstructing the muscle component of cultivated meat, it has become increasingly clear that muscle tissue alone cannot replicate the complete sensory and functional qualities of conventional meat. Fat tissue, a critical determinant of flavor, aroma, texture, and mouthfeel, poses a significant challenge for *in vitro* reproduction [4–6]. Achieving the desired organoleptic properties necessitates precise engineering of the fat component, which remains a major focus of current research efforts.

The successful differentiation of stem cells into adipocytes involves a complex interplay between the cellular microenvironment and biochemical signals. Adipogenesis requires a carefully orchestrated 3D culture environment that mimics native tissue conditions. Factors such as cell-cell interactions, extracellular matrix (ECM) composition, substrate stiffness, and the presence of specific biochemical inducers are essential to drive the commitment, differentiation, and maturation of stem cells into lipid-storing adipocytes [7–10]. Notably, adipogenic differentiation relies heavily on the activation of the peroxisome proliferator-activated receptor gamma (PPARγ) pathway, a key regulatory mechanism for adipocyte development [11,12]. Therefore, the composition of the culture medium, particularly the selection and concentration of fatty acids, will directly influence the final lipid profile, which in turn determines the taste, aroma, and texture of the reconstructed fat tissue [13–15]. Additionally, the use of edible biomaterials such as gelatin and alginate can allow fine control over matrix stiffness and porosity, which are critical parameters influencing adipocyte maturation and lipid droplet formation [16–18] for enhanced differentiation quality and physical realism. These structural advances are essential for also reproducing the meat textural properties, such as tenderness and juiciness, that drive consumer preference.

Despite the significant progress achieved in generating *in vitro* adipose tissues for food applications, major limitations remain, particularly in producing edible bovine fat that exhibits organoleptic properties comparable to native tissue. Many studies still rely on murine models such as 3T3-L1 cells [19,20], while most other species-specific systems have used porcine, caprine or chicken cells as models [4,16,19,21–30]. These approaches, however, cannot reproduce the bovine-specific compound profiles that are critical for authentic flavor reproduction. When bovine-derived cells have been explored, including fibro-adipogenic progenitors [31], mesenchymal stem cells [32,33], umbilical cord stem cells [34], fibroblasts [35], and adipose-derived stem cells [36–38], some studies have evaluated only adipogenic differentiation efficiency [32,34,35], while most organoleptic assessments have been limited to basic lipid profiling [31,33,37,38]. Only a few have extended analyses to include flavor compound profiling [36] or texture assessments that capture the dynamic mouthfeel of the tissue [35]. The most critical limitation, however, lies in the adipogenic differentiation media employed across almost all published protocols. These typically depend on non-edible pharmacological inducers such as insulin, dexamethasone, rosiglitazone or other thiazolidinediones, IBMX, indomethacin, or doxycycline [4,16,19–22,25–28,31–38]. Such compounds are unsuitable for food-grade cultivated fat production due to safety and regulatory restrictions. Encouragingly, a few recent studies have begun to explore food-compatible adipogenic supplements, including plant oils such as olive, soybean, peanut, sunflower, or tea seed oil, and Intralipid emulsions [4,19,21–24,34,37]. These efforts represent important first steps toward the development of edible, regulatory-compliant systems for cultivated bovine fat production, though their efficacy in replicating native flavor and texture remains limited.

In our previous study [15], we established a food-grade differentiation protocol for bovine adipose-derived stem cells (bADSCs), demonstrating that it is possible to achieve a lipid profile closely resembling that of natural bovine fat. This work provided a foundational basis for developing edible, bovine-specific adipose tissues under biocompatible culture conditions. However, several aspects related to lipid yield optimization, scaffold selection, and sensory-relevant characterization remained to be addressed. These parameters are critical for replicating not only the chemical composition but also the textural and organoleptic features necessary for consumer acceptance.

To address these remaining challenges, the present study focuses on establishing a fully food-grade platform for cultivated bovine fat fabrication. We evaluate the effects of food-grade scaffolds and culture media on adipogenic differentiation, lipid accumulation, and tissue structure. The resulting engineered fat is characterized through quantitative analyses of fatty acid composition, lipid yield, and physical texture, and benchmarked against natural bovine fat to assess sensory-relevant equivalence.

Beyond improving culture performance, this study also aims to align cultivated fat production with food safety and consumer acceptance standards. The exclusive use of edible components ensures compatibility with regulatory frameworks for food products, while detailed lipid and structural characterization strengthens confidence in the authenticity and quality of the final tissue. By integrating these scientific and translational perspectives, this work provides a comprehensive foundation for developing organoleptically realistic and commercially viable bovine cultivated fat.

## Materials and Methods

### Materials and products

Fetal bovine serum (FBS, 35010CV) was purchased from Corning (NY, USA). Hexane (17922-94), methanol (21915-35), NaOH (31511-05), NaCl (31320-34) for fatty acid analysis, DMEM (08458-16), Antibiotics-Antimycotics (02892-54) and PBS (07269-84) were sourced from Nacalai Tesque (Kyoto, Japan). Bovine fibrinogen (F8630), bovine thrombin (T4648), gelatin (G1890), agarose (A9045), lactalbumin hydrolysate (Sigma, 61302), FAME Mix and BF3/MeOH/MeOH solution (B1252) were purchased from Sigma Aldrich (Saint-Louis, USA). CaCl2 (58001-17) came from Kanto Chemical (Tokyo, Japan), food-grade CaCl2 from Shiraimatsu Pharmaceutical Co. Ltd. (Shiga, Japan). EZ-BindShut 96 well plates, lid round bottom were sourced from Iwaki (AGC TECHNO GLASS CO.,LTD. Shizuoka, Japan). Perasan MP2-J was from Enviro Tech Japan Co., Ltd. (Tokyo, Japan). Albumin from Bovine Serum Fatty Acid Free (011-15144), Trypsin (209-19182), and PFA 4% (16310245) were from Wako (FUJIFILM, Japan). iScript™ cDNA Synthesis Kit (1708890) was from BioRad, Applied Biosystems™ TaqMan™ Fast Advanced Master Mix (4444557) and Hoechst 33324 (H3570) from Thermo Fisher Scientific (Waltham, USA). Nile Red (N0659), gum Arabic (A3553) and carrageenan (C3313) were sourced from Tokyo Chemical Industry Co., Ltd. (Japan). Hyaluronan (GS311 HyStem®, Thiol-Modified Hyaluronic Acid Kit) was from Cellink (BICO, Göteborg, Sweden). Food-grade medium I-MEM was purchased from IntegriCulture Inc. (Fujisawa, Japan), food-grade oleic acid from NOF Corporation (Tokyo, Japan) and food-grade sodium alginate from Marugo Corporation (Okayama, Japan). Distilled water used for GC–MS analysis was pre-washed with hexane (Kanto Chemical). Whey proteins (WP) (Milei80, MILEI GmbH, Leutkirch, Germany) were kindly provided by Morinaga Milk Industry Co., Ltd. (Kanagawa, Japan).

### Livestock tissues used and isolation, purification of bovine adipose-derived stem cells (ADSCs)

Bovine tissues were obtained from commercial Japanese Black cattle fresh beef isolated after slaughter (Ariake Beef Plant of 544 SANKYOMEAT INC., Japan). The tissues were kept at 0-4 °C and carried to the laboratory after health inspection of the carcass. On the next day, fat surrounding shintama (Knuckle) muscle were separated from the muscle tissues and used for cell isolation from each individual animal, following previously published method with slight modifications [15]. Briefly, to be handled in accordance with the “Food Sanitation Act” ), the tissue samples were first disinfected using Perasan MP2-J (Enviro Tech Japan, Tokyo, Japan) at 0.18 % (w/v) as peracetic acid and the digestion was performed using edible trypsin. Isolated bADSC were cultivated in high-glucose IMEM (IntegriCulture Minimum Essential Media, Japan) containing 10% of edible bovine serum. After 3 days of attachment and expansion, the cells were washed 3 times with PBS to remove all the remaining tissue debris and the medium was renewed every 2-3 days until 70% of confluence where the cells were passaged. After one passage, the remaining adherent cells were considered to be bADSC.

The cells were cryopreserved in Bambanker DMSO Free (NIPPON Genetics, Tokyo, Japan). In this study, one lot means a single bovine donor.

The edible bovine blood serum was prepared by modifying the edible grade bovine plasma derived from Proliant (New Zealand). CaCl2 was added to the bovine plasma at a final concentration of 20 mM and incubated to solidify overnight at 37 °C, then filtered through a 0.22 µm strainer. The obtained liquid was used as the bovine blood serum.

### 3D bADSC culture

To evaluate different biocompatible scaffolds for bovine adipogenesis, bovine ADSCs were first detached using trypsin and centrifuged (1,970 g, 1 min). The cell pellets were mixed with sterilized edible biomaterial solutions, previously autoclaved (121 °C, 20 min) and stored at −20 °C until use, to obtain a final concentration of 5 × 10 cells/mL. The formulations tested included: alginate (1% final concentration, prepared in PBS and crosslinked by dropping on 100 mM CaCl solution in PBS); gum arabic/alginate (1%/0.5% final concentrations, prepared in PBS, crosslinked by dropping on 100 mM CaCl solution); gellan gum (1% final concentration, prepared in Milli-Q water, crosslinked with 0.1% CaCl , 2:1 ratio); κ-carrageenan (1.5% final concentration, prepared in Milli-Q water, crosslinked by mixing with 0.1% CaCl ); agarose (1.5% final concentration, low-melting-point, prepared in PBS and gelled by cooling to room temperature); hyaluronic acid (0.8% final concentration, prepared using the HyStem® thiol-modified HA kit; Advanced Biomatrix); and fibrin (6 mg/mL fibrinogen with 3 U/mL thrombin final concentrations). For each condition, 10 µL cell–biomaterial mixtures balls tissues were prepared and allowed to gel for 5 min. The resulting 3D constructs were washed three times in PBS and transferred to low-attachment 96-well plates (EZ-BindShut ) containing 300 µL of IMEM medium supplemented with 10% bovine blood serum. After 2 days of proliferation to enable enough cell–cell contact, the medium was replaced with IMEM supplemented with 10% bovine blood serum and oleic acid at various concentrations for 7 to 28 days of differentiation, with medium renewal every 2–3 days.

### BSA, Lactalbumin and WP-oleic acid complexes

BSA/Lactalbumin/WP–oleic acid (OA) complexes were prepared at a molar ratio of 1:6 (BSA/Lactalbumin/WP:oleic acid), with a final stock concentration of 10 mM oleic acid. Briefly, 150 mM sodium chloride solution was prepared in Milli-Q water. BSA/Lactalbumin/WP (final concentration 1.6 mM) were dissolved in this solution and incubated at 37 °C for 20 min to ensure complete solubilization, followed by sterile filtration through a 0.22 µm filter. Separately, oleic acid was also sterilized by filtration (0.22 µm) and added to the BSA/Lactalbumin/WP solution at a final concentration of 10 mM. The mixture was vigorously manually agitated for 20 s and incubated in a 37 °C water bath for 1 h to allow complex formation. For bovine adipogenic differentiation medium, the stock solution was then diluted to a final oleic acid concentration of 1.5 mM in the culture medium.

Concerning the WP, we used the Milei80 (MILEI GmbH, Leutkirch, Germany) which is an edible WP protein concentrate with:

- Protein: 80.0% or more per solids
- Lactose: 6.5% (max 8.5%)
- Ash: 3.0% (max 4.0%)
- Fat: 5.0% (max 7.0%)
- Moisture: (max 5.0%)

including β-lactoglobulin (52%), α-lactalbumin (14%), bovine serum albumin (9%), and immunoglobulins (7%).

### 3D bioprinting fibers and culture

Fibers printing was conducted in the same way as already published, with slight modifications, in gelatin supporting bath [39]. Briefly, granular edible gelatin was produced by preparing 4.5 wt% gelatin in IMEM containing 10 % bovine blood serum and 20 mM CaCl2, putting it at 4 overnight for gelation, adding the same volume of IMEM-CaCl2 to the gelatin gel, grinding it with a rotor-stator homogenizer for 2 min at 30,000 rpm, centrifuging at 2,837 g for 3 min, and removing the supernatant. The bioink was prepared by mixing 5x10^6^ cells/mL of bADSCs in the autoclaved 1% alginate. After filling the prepared Gelatin-CaCl2 supporting bath in a 24 well plate and loading the syringe containing the prepared bioink onto the dispenser instrument (Musashi, Shotmaster 200DS), cell printing was conducted inside the supporting bath. All parts, such as syringes, nozzles, and containers used for cell printing, were first sterilized by autoclave. The nozzle gauge, moving speed, and dispensing speed was 14G, 1 mm/s, and 2 uL/s, respectively. The printed structures inside the supporting baths were then incubated at room temperature for 5 min to ensure full gelation and the gelatin bath was dissolved by incubation at 37 for 1 h. The liquid gelatin was then removed and the obtained cell fibers were cultivated for 2-3 days and then replaced with the differentiation medium.

### Compression Testing

Compression tests were performed at room temperature using the EZ-TEST instrument (AGS-X, Shimadzu, Japan) equipped with a 20 N load cell and an 8 mm diameter plate probe. Hydrogels (200 μL) with varying concentrations of alginate were cast in 24-well plate inserts. After gelation in CaCl2 solution, the samples were transferred to the instrument for analysis. Each hydrogel sample’s diameter and height were measured with a vernier caliper prior to testing. Samples were compressed at a constant rate of 20.0 μm/min. The deformation was recorded continuously to generate stress-strain curves, from which the elastic modulus was determined. The modulus was calculated within the 10%–20% strain range of the sample thickness using TRAPEZIUM LITE X software (Version 1.5.8, Shimadzu, Japan).

### Rheological Characterization

The rheological properties of the hydrogels were evaluated using a rotational rheometer (MCR302, Anton Paar, Graz, Austria). Hydrogels (250 μL) were cast in 6-well plate inserts to get 25 mm diameter and 5 mm thickness disks. After gelation in CaCl2 solution, the samples were transferred to the rheometer for analysis. *In vivo* samples were also cut at the same dimensions before measurement. Frequency sweep measurements were conducted from 0.1 to 10 rad/s at a fixed strain amplitude of 1.0%, with an initial axial force of 0.2 N and at a temperature of 25°C. The storage modulus (G′), representing the elastic or solid-like behavior, and the loss modulus (G″), representing the viscous or liquid-like behavior, were recorded over a frequency range of 0.1 to 10 Hz. For each alginate concentration as well as for *in vitro* and *in vivo* samples, frequency-dependent moduli were plotted to assess the material’s viscoelastic behavior.

### Gene expression

Gene expression was analyzed using real-time quantitative polymerase chain reaction (RT-qPCR). Adipose ball tissues in fibrin and in alginate gels, at day 7 of differentiation, were washed in PBS and total RNA extraction was carried out using the PicoPure^TM^ RNA Isolation Kit (Arcturus, ThermoFisher Scientific, USA), with the DNAse step, following the manufacturer’s instructions. Samples’ RNA content was quantified with the NanodropTM spectrometer (N1000, Thermo Fisher Scientific, Waltham, USA). For RT-qPCR, the RNA samples were first submitted to reverse transcription into cDNA using iSCRIPT cDNA synthesis kit (Bio-Rad, Hercules, USA), before being amplified using Taqman probes and reagents (Taqman Fast Advanced Mix, Taqman gene expression assays (FAM): ITGA5 (Assay ID: Bt04299016_m1), ITGA6 (Assay ID: Bt03276105_m1), VIM (Assay ID: Bt03244344_m1), ACTB (Assay ID: Bt03279174_g1), TUBA1A (Assay ID: Bt04299043_g1), FABP4 (Assay ID: Bt03213820_m1), PPARG (Assay ID: Bt03217547_m1) and PPIA (Assay ID: Bt03224615_g1), Thermo Fisher Scientific, Waltham, USA). The cDNA synthesis and RT-qPCR reactions were conducted using the StepOnePlus Real-Time PCR System (Thermo Fisher Scientific, Waltham, USA) and the gene expression was normalized by PPIA as the housekeeping gene.

### Lipogenesis analysis

Following adipogenesis, the cells were fixed in 1% paraformaldehyde prepared in 50 mM CaCl2 in PBS at 4 °C overnight and 100 ng/mL Nile Red was applied for lipid staining at room temperature for 30 min, counterstained with Hoechst for nuclei. For the calculation of lipid production from bADSC, the 3D tissues’ Z-stack images were taken with the same exposure time, brightness, and contrast for all conditions. Then, the Maximum Intensity Projection (MIP) of Z-slice’s images in Nile Red and Hoechst of each 3D tissue was performed. For this, a Confocal Quantitative Image Cytometer CQ1 (Yokogawa, Tokyo, Japan) was used, with the following channels: Nile Red using the 561 nm laser (power 20%), Ex: 617∼673 nm, exposure time 500 ms, and Hoechst using the 405 nm laser (power 100%), Ex: 447∼460 nm, exposure time 500 ms. The total fluorescence intensity was finally calculated from the total Nile Red’s intensity of the MIP images divided by the total Hoechst intensity in each 3D tissue using ImageJ software (version 1.53j, NIH, USA).

For the lipid vesicle size assessment using Nile Red and Hoechst staining projection images, MRI_Lipid Droplets plugin tool was used (http://dev.mri.cnrs.fr/projects/imagej-macros/wiki/Lipid_Droplets_Tool) [40] in the ImageJ Software (NIH) with slight modification to analyze either the red or the blue channel. On 20x magnification images, the unilocular lipid vesicles with a diameter above 35 µm were selected, being considered as enough mature, and the number of mature unilocular vesicles was divided by the total number of nuclei per image (for each condition, n=6-8 different pictures were counted from 3-4 different samples).

For the quantitative triglycerides’ analysis, 5 balls for each sample were homogenized in NP40 Substitute Assay Reagent buffer solution and then centrifuged for 10 min at 10,000g in 4 °C. Triglycerides concentrations were then measured on the supernatant following the protocol’s kit (10009582, Cayman Chemical Company, Michigan, USA). The triglycerides content was finally normalized by the DNA amount, measured on the same supernatant lysates using the Qubit™ dsDNA HS Assay Kit (ThermoFisher, USA).

### Fatty acids composition assessment

Samples shown in Figure 4 were analyzed at NISSIN FOODS Holdings, Global Food Safety Institute, following their established GC–FID protocol for fatty acid methyl ester (FAME) profiling as previously described [15]. In brief, lipids extracted from cultivated bovine fat and bovine cheek intramuscular fat controls were saponified, methylated, and analyzed on a Shimadzu GC 2010 Plus gas chromatograph equipped with a DB 23 capillary column (Agilent Technologies). Fatty acids were identified by comparison with certified standard mixtures (fatty acid methyl ester (FAME) Mix, included 37 FAMEs), and methyl heneicosanoate (C21:0) was used as an internal standard.

Samples presented in Figure 7 were analyzed by Shimadzu Techno Research, Inc., using their validated GC–MS method for quantitative fatty acid composition. Frozen tissue samples were subjected to methanolysis and analyzed on a GCMS QP^TM^ 2020 system (Shimadzu) with a TC 70 capillary column (60 m × 0.25 mm i.d., 0.25 μm film thickness; GL Sciences, J2102K0). Calibration and quantification were performed using authentic FAME standards with palmitic acid d as an internal standard with linearity confirmed (p > 0.9990).

### Statistical analysis

Statistical analysis was performed using EzAnova (version 0.98, University of South Carolina, Columbia, SC, USA) software. The detail of the number of n corresponding to the number of independent experiments using isolated bovine primary cells from different bovine donors, or independent samples, is displayed in the captions. For two-way ANOVA, time set was applied as “paired or repeated measures” and the treatment as classic analysis or “unpaired” which led to a pairwise comparison, with the Tukey’s HSD post hoc test for the multiple comparisons. Error bars represent SD. *p* values <0.05 were considered to be statistically significant.

### Ethical approval

Ethical approval was not required for this study since the bovine ADSC were isolated from commercial fresh beef obtained after slaughter and sent to the laboratory.

### Data availability

All data supporting the key findings of this study are available from the corresponding authors upon reasonable request.

### Authors contribution statement

Conceptualization: F.L., S.T., M.M. ; Data curation: F.L., A.C, M.F. ; Formal analysis: F.L., A.C, MF ; Funding acquisition: S.T., S.K., M.M.; Investigation: F.L., M.F., S.T., M.M.; Methodology: F.L., M.F., S.T., M.M.; Project administration: S.T., S.K., M.M.; Resources: S.T., S.K., M.M. ; Validation: F.L., M.F., S.T., S.K., M.M.; Writing - original draft: F.L., M.F., T.N. ; Writing - review & editing: F.L., M.F., T.N., S.T., M.M.

## Results and Discussion

### Determining the most suitable material for bovine adipogenesis

To enhance current strategies for bovine adipogenesis in cultivated meat production, a comparative analysis of various edible biomaterials was first conducted to identify a matrix capable of promoting robust lipid accumulation from bADSC (Figure 1). For materials requiring CaCl -induced crosslinking, droplets containing bADSC) were deposited into a CaCl solution to form spherical tissue constructs of approximately 10 µL in volume (Figure 1A-B). In the case of self-gelling materials, such as hyaluronic acid, the gelation initiator was incorporated directly into the 10 µL constructs. Following 7 days of adipogenic differentiation in medium supplemented with 1.5 mM of oleic acid, alginate-based constructs already exhibited prominent unilocular mature adipocytes, as visualized by phase-contrast microscopy (Figure 1C) and Oil Red O staining (Figure 1D). To more readily qualitatively assess lipid accumulation, cells were then stained with Nile Red, a fluorescent dye specific for neutral lipids, and counterstained with Hoechst to visualize nuclei. Nile Red staining also revealed a high proportion of mature adipocytes within the alginate constructs (Figure 1E). In comparison, all other tested biomaterials exhibited lower adipogenic efficiency, with the exception of the alginate–gum arabic composite. Notably, gum arabic alone was unable to form stable gels, highlighting the importance of alginate in maintaining structural integrity (Figure 1F–J). Based on these findings, alginate was selected for subsequent experiments.

**Figure 1.**
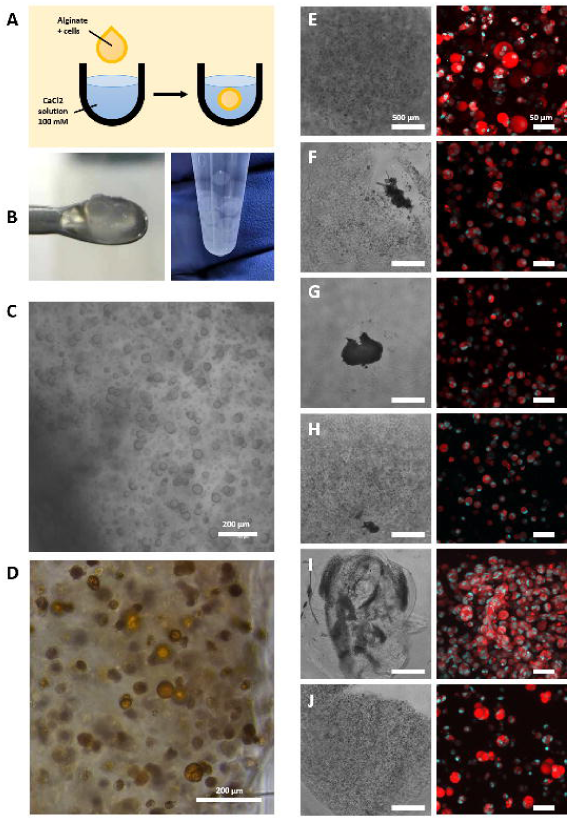
Screening of edible scaffolds for bovine adipogenesis. (A) Schematic of the 3D culture setup for seeding of 10 μL droplets of bADSCs encapsulated in CaCl -crosslinked hydrogels matrices. (B) Representative images of ball tissue constructs, here formed with alginate 1%. (C) Phase-contrast microscopy representative images of bovine adipose constructs after 7 days of differentiation in alginate gels conditions. (D) Oil Red O staining representative images of bovine adipose constructs after 7 days of differentiation in alginate. (E) Confocal representative images of 3D bovine adipose constructs after 7 days of differentiation in alginate, stained with Nile Red (lipids) and Hoechst (nuclei). (F–J) Representative images of lipid accumulation by Nile Red/Hoechst fluorescence in other biomaterials tested (gellan gum, -carrageenan, agarose, hyaluronic acid, and gum arabic + alginate, in order).

### Optimization of the alginate culture condition for enhanced bovine adipogenesis

Next, the concentration of alginate was optimized to establish the optimal mechanical properties for supporting bovine adipogenesis (Figure 2). Alginate concentrations ranging from 0.25% to 10% were evaluated. Constructs formed with 0.25% alginate failed to maintain structural integrity until the end of the 7 days adipogenic differentiation culture, while those with 5% and 10% alginate exhibited markedly reduced lipid accumulation and adipocyte maturation (Figure 2A). Quantification of intracellular triglyceride content confirmed these results, revealing a significant 48% and 68% reduction in lipid content in 5% and 10% alginate constructs, respectively, compared to 1% alginate condition (Figure 2B). DNA quantification further indicated impaired cell proliferation in the denser gels, with a 27% and 55% decrease in DNA content in 5% and 10% alginate, respectively, relative to 1% (Figure 2C). These data suggest that high alginate concentrations can hinder cellular aggregation, which is critical for efficient adipogenic differentiation. Aggregation of preadipocytes into dense 3D clusters is a well-documented prerequisite for initiating adipogenesis, as it facilitates enhanced cell–cell contact and localized accumulation of adipogenic signals such as PPARγ ligands [41]. Additionally, dense ECM environments, such as those formed by high-concentration alginate, can impose physical constraints that not only limit cellular proliferation but also inhibit the morphological rounding required for terminal adipocyte maturation [42,43]. The superior adipogenesis observed in 1% alginate thus reflects both structural permissiveness and mechanical compatibility. Specifically, stiffer matrices have been shown to suppress adipogenesis by interfering with the mechanotransduction pathways necessary for adipogenic gene activation, including YAP/TAZ signaling and cytoskeletal remodeling [44]. Our findings align with these studies, suggesting that lower alginate concentrations provide a mechanically permissive environment that supports cellular aggregation and promotes adipogenic commitment.

**Figure 2.**
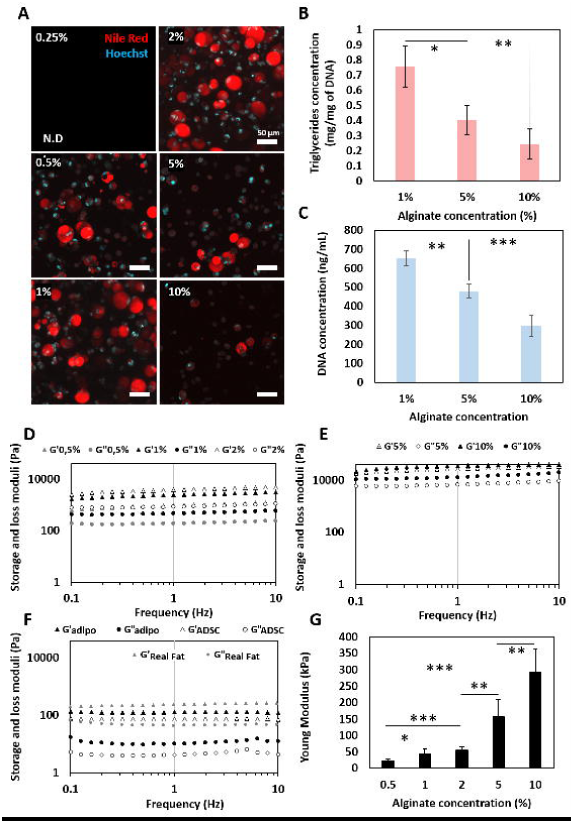
Optimization of alginate scaffold concentration for adipogenesis and mechanical fidelity. (A) Representative Nile Red and Hoechst fluorescence images of bADSC encapsulated in alginate constructs of varying concentrations (0.25–10%) followed by adipogenic differentiation for 7 days. (B) Quantification of intracellular triglyceride content normalized to DNA, showing lipid accumulation in response to varying alginate concentrations (1–10%) . N = 3 independent donors replicates. (C) DNA content analysis of the differentiated constructs, in response to varying alginate oncentrations (1–10%). N = 3 independent donors replicates. (D-E) Storage modulus (G′) and loss modulus (G″) of cell-free alginate gels assessed by frequency sweep rheometry in varying alginate concentrations (0.5–10%) . (F) Rheological comparison of differentiated adipose tissues (“Adipo”), undifferentiated ADSC-laden gels (“ADSC”), and native bovine fat (“Real Fat”). (G) Young’s modulus measured by compression testing on cell-free alginate gels with varying alginate concentrations. n = 3–6 replicates. Error bars represent mean ± SD. Asterisk (*) denotes significant statistical enrichments (\**p*<0.05, ** *p*<0.01 and *** *p*<0.001).

To further evaluate the suitability of the alginate-based hydrogels as fat mimetics in cultivated meat fat, we therefore also analyzed their mechanical and viscoelastic properties both without cells and immediately after gelation (Figure 2D,E) as well as after cell culture under adipogenic or proliferation conditions, and *in vivo* bovine fat (Figure 2F, “adipo” meaning differentiated adipocytes, “ADSC” meaning proliferating undifferentiated ADSC, “Real Fat” meaning *in vivo* bovine fat tissues). Frequency sweep tests of cell-free alginate gels revealed that all formulations displayed a predominantly elastic behavior, with storage modulus (G′) consistently exceeding loss modulus (G″). At low alginate concentrations (0.5–2%), the hydrogels were relatively soft, with G′ values ranging from ∼600 Pa to ∼5000 Pa, corresponding to estimated Young’s moduli between ∼1.8 and ∼15 kPa (based on the approximation E≈3G′). These values align well with the texture of raw or lightly cooked animal fat, such as intramuscular fat in Wagyu beef [45,46], which contributes to the signature “melt-in-the-mouth” sensation during eating [47,48]. Importantly, such soft textures enhance flavor release by enabling rapid fat breakdown and lipid dissolution at oral temperatures, mimicking the behavior of native fat globules [49,50]. This is particularly relevant for replicating the creamy mouthfeel and juiciness associated with premium marbled meats. In contrast, at higher alginate concentrations (5–10%), G′ values exceeded 10,000 Pa (estimated E ∼30–60 kPa), leading to considerably firmer gels with a rubberier consistency that do not replicate the delicate mouthfeel of premium marbled fat [47,48].

After 7 days of differentiation culture, the rheological profiles of alginate samples containing either differentiated bovine adipocytes or undifferentiated bADSCs (Figure 2F, cell-free sample controls with same culture duration in Supplementary Figure 1) were also characterized. Adipocyte-laden hydrogels (adipo) showed higher viscoelastic moduli than their ADSC counterparts (G′ ∼130 Pa vs. ∼75 Pa at 1 Hz), indicating matrix remodeling and lipid accumulation during adipogenesis. These G′ values correspond to estimated Young’s moduli of ∼450 Pa and ∼300 Pa, respectively, both well within the mechanical range reported for natural adipose tissue in raw meat [46] and also measured on our isolated *in vivo* real fat (from shintama muscle), supporting its suitability for cultivated meat applications. Such soft gels should mimic the spreadability and thermal melting of real fat during cooking, supporting oil release and flavor blooming under mild heating conditions (e.g., <100°C). From a sensory perspective, their low stiffness enhances oral disintegration, helping to recreate the lubrication and smooth texture consumers expect from animal-derived fat with desirable organoleptic features such as mouthfeel, juiciness, and flavor release during cooking and mastication.

On the other hand, the compression testing (Figure 2G) performed on cell-free alginate gels at different concentrations showed a pronounced increase in Young’s modulus with alginate concentration, from ∼25 kPa at 0.5% to 300 kPa at 10%. These values, though considerably higher than rheological estimates, reflect the gels’ bulk mechanical resistance under uniaxial stress. This difference may result from structural factors such as gel network density, water expulsion under load, and increased ionic crosslinking at higher alginate contents. For food applications, this elevated compression modulus suggests improved handling, slicing, and shape retention, which are critical for product processing and presentation. However, formulations exceeding 50–100 kPa may risk over-hardening, leading to a dry or rubbery mouthfeel not typically associated with palatable fat [49]. Thus, for applications prioritizing eating quality, lower concentrations (0.5–2% alginate) appear more favorable.

All these properties are critical for replicating the sensory and functional qualities of real meat fat, such as softness during mastication, efficient flavor release, and sufficient structural integrity for cooking and handling. While the 0.5% alginate gel most closely matched the mechanical softness of native fat, we selected 1% alginate for the following experiments as it offered a more practical balance between physiological texture and handling stability during manipulation, cell culture, and downstream processing.

### Adipogenic, cytoskeletal and integrin remodeling gene expression during bADSC differentiation

To further characterize and understand the induced adipogenic differentiation in alginate compared to less suitable biomaterials like our previous fibrin-based method [15], qPCR analysis of key gene markers was performed (Figure 3A-C). Two well-established adipogenesis-related genes were assessed: PPARγ (Peroxisome proliferator-activated receptor gamma), an early adipogenic marker, and FABP4 (Fatty acid-binding protein 4), a marker of terminal adipocyte maturation. PPARγ plays a critical role in initiating the adipogenic transcriptional cascade in ADSC, activating downstream genes that promote lipid uptake and storage [51,52]. FABP4, also known as aP2, binds and transports long-chain fatty acids and is essential for lipid droplet formation and intracellular lipid homeostasis [53,54]. The bovine adipocytes differentiated for 7 days in 1% alginate exhibited significant 3-fold increase in PPARγ expression and 32-fold increase in FABP4 expression relative to fibrin culture condition, indicating enhanced adipogenic commitment and maturation. These findings are consistent with previous studies that show high FABP4 expression is a hallmark of mature unilocular adipocytes, while elevated PPARγ expression is necessary to drive preadipocyte commitment into the adipogenic lineage and both are specifically found enhanced in 3D culture condition [8].

**Figure 3.**
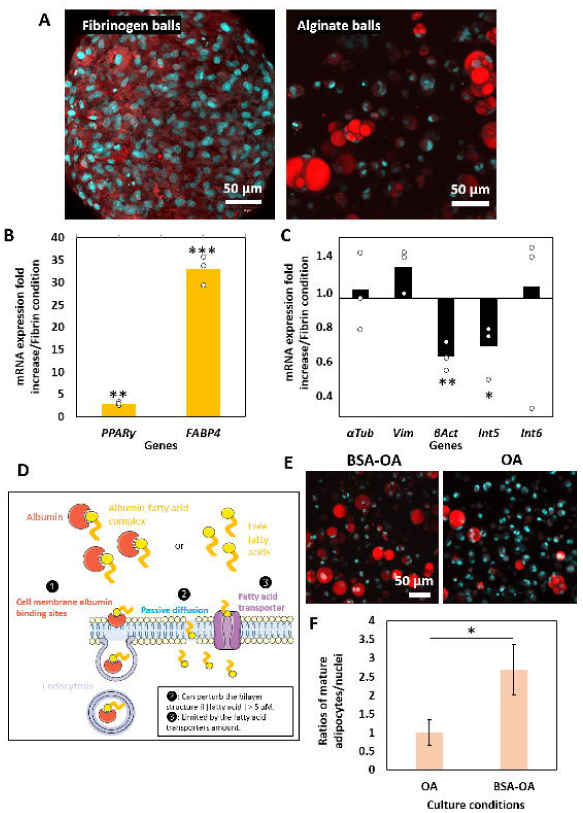
Fluorescence imaging of lipid accumulation and gene expression analysis in bADSCs cultured in alginate versus fibrin, along with the effect of BSA-mediated oleic acid delivery. (A) Confocal representative images of 3D bovine adipose constructs after 7 days of differentiation in fibrin or 1% alginate gels conditions, stained with Nile Red and Hoechst. (B-C) Relative gene expression of adipogenic markers (FABP4, PPARG), cytoskeletal markers (VIM, αTUB, βACT), and integrin subunits (ITGA5, ITGA6) in alginate versus fibrin constructs, normalized to PPIA, after 7 days of differentiation. N = 3 independent donors replicates. (D) Schematic representation of possible cellular fatty acid transport in bovine ADSC. (E) Confocal representative images of 3D alginate constructs after 7 days of differentiation with oleic acid alone or BSA-complexed oleic acid, stained with Nile Red and Hoechst. (F) Quantification of mature adipocytes ratio (number or lipid vesicles with diameter above 35 µm^2^) normalized by the total number of nuclei, comparing unbound and BSA-complexed oleic acid conditions. n = 3 biological replicates. Error bars represent mean ± SD. Asterisk (*) denotes significant statistical enrichments (\**p*<0.05, ** *p*<0.01 and *** *p*<0.001).

**Figure 4.**
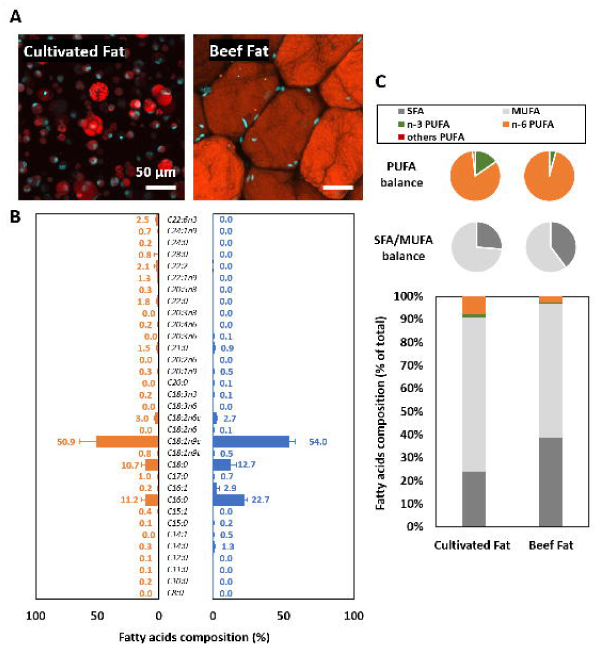
Comparison of lipid morphology and fatty acid composition between engineered cultured fat and native beef fat. (A) Confocal microscopy images of cultured fat (left) and native beef fat (right) stained with Nile Red and Hoechst. (B) Fatty acid composition of cultured fat (orange) and native beef fat (blue), shown as percentage of total fatty acids. (C) Summary of fatty acid classes in cultured and native fat: PUFA balance (top), SFA/MUFA balance (middle), and total fatty acid class distribution (bottom). N = 3 independent donors replicates for native beef fat and N = 6 independent donors replicates for cultured fat samples. Error bars represent mean ± SD.

Since the cells showed different morphology in fibrin and alginate culture conditions, we also assessed the expression of cytoskeletal genes associated with adipogenesis. Vimentin (Vim) was modestly upregulated (1.3-fold) in alginate compared to fibrin, which aligns with previous studies showing its dynamic reorganization during adipogenesis, where it functions as a structural scaffold for lipid droplet formation, mediates their trafficking and clustering, and correlates with increased intracellular triglyceride accumulation [55,56]. α-Tubulin (αTub) also increased slightly (1.1-fold), supporting the notion of active microtubule reorganization during differentiation. Conversely, β-actin (βAct) was significantly downregulated by 0.6-fold, consistent with the cytoskeletal reconfiguration required for adipocyte rounding and lipid accumulation. These results reflect the coordinated cytoskeletal remodeling required for adipocyte differentiation, consistent with previous reports of actin depolymerization and microtubule reorganization during lipid droplet accumulation and preadipocytes transition to mature adipocytes [57,58].

In addition, changes in integrin expression were also investigated, given their known role in extracellular matrix (ECM) interactions during adipogenesis. Integrin α5 (Int5) was significantly downregulated (0.7-fold), while Integrin α6 (Int6) was slightly upregulated (1.1-fold). These trends are consistent with a transition from fibronectin-to laminin-mediated ECM interactions, as described by Uetaki et al., where decreased Int5 facilitates actin remodeling and increased Int6 supports laminin engagement during adipocyte maturation [59].

All these findings suggest that even within alginate, which is a non-adhesive alginate scaffold, bADSCs can create a conducive microenvironment, possibly through ECM secretion, enabling integrin-mediated signaling pathways that promote adipocyte maturation.

### The importance of albumin complexation for oleic acid adipogenic induction

We have previously showed that incorporation of oleic acid into the culture medium significantly enhanced adipogenic differentiation of bADSCs in a similar way than *in vivo* [15]. To further potentiate this effect, we evaluated the impact of complexing oleic acid with albumin, using first Bovine Serum Albumin (BSA), which acts as a carrier protein. BSAcan bind to fatty acids, increasing their solubility in aqueous environments and facilitating their transport to the cell membrane, mimicking physiological conditions where albumin shuttles non-esterified fatty acids through the bloodstream. This complexation not only enhances cellular uptake, potentially via receptor-mediated transport or diffusion, but also minimizes lipotoxicity associated with high concentrations of unbound fatty acids [60,61] (Figure 3D). To assess the effect of BSA-complexed oleic acid, we performed quantitative analysis of lipid accumulation from Nile Red staining (Figure 3E-F). Cultures treated with the BSA-oleic acid complex exhibited a significant 2.7-fold increase in lipid storage compared to those treated with oleic acid alone, indicating a more efficient and robust adipogenic response.

### Organoleptic assessment of bovine adipocytes alginate tissues by lipid composition

To evaluate how closely our engineered adipose tissue mimics native bovine fat, we compared the fatty acid composition and total lipid content of in *vitro* differentiated bovine ADSC during 7 days to native beef fat (Figure 4). Microscopy images (Figure 4A) revealed morphological differences in lipid droplet structure: native bovine fat exhibited large, tightly packed unilocular droplets, while the cultivated fat contained smaller, more heterogeneous droplets dispersed within the alginate matrix. These observations were consistent with lipid quantification, which showed approximately 4 mg/g of fatty acids in cultivated fat versus ∼160 mg/g in native tissue, meaning a ∼36-fold difference. Although total lipid content in our cultivated fat remains lower than *in vivo* fat, this result already provides a valuable benchmark for lipid productivity in cultivated adipose tissue because previous reports focused only on relative fatty acid profiles or qualitative lipid staining.

Despite this reduced total lipid content, the fatty acid composition of the cultivated fat closely approximated that of native bovine adipose tissue (Figure 4B–C). During the adipogenic differentiation of the bADSCs, there is normally a notable shift in the fatty acid composition within the cells. Specifically, the proportion of saturated fatty acids (SFAs) decreases, while monounsaturated fatty acids (MUFAs), such as oleic acid (C18:1n9c), increases. This change is attributed to active de novo lipogenesis and desaturation processes during adipogenesis [62]. Oleic acid (C18:1n9c), the dominant monounsaturated fatty acid (MUFA) in Wagyu beef, was indeed found the most abundant fatty acid in both samples, comprising 50.9% in cultivated fat and 54.0% in native beef. This alignment is important given oleic acid’s contribution to fat softness, low melting point, and rich umami flavor [63–68].

Major saturated fatty acids (SFA) such as stearic acid (C18:0) and palmitic acid (C16:0) were also detected in cultivated fat at 11.2% and 10.7%, respectively, values that are comparable to those in native fat (12.7% and 22.7%). The stearic-to-palmitic acid ratio in cultivated fat appears favorable, as stearic acid is considered metabolically neutral in humans, unlike palmitic acid which has been associated with lipotoxicity [69]. Palmitoleic acid (C16:1), a minor MUFA with bioactive properties, was present at 0.8% in cultivated samples and 2.9% in native tissue. These values result also in a favorable SFA:MUFA balance, consistent with the high MUFA:SFA ratio observed in Wagyu beef (ranging from 2.08 to 2.57 depending on grade) [63–68].

Interestingly, “other” fatty acids, including minor polyunsaturated fatty acids (PUFAs) and atypical chain lengths, comprised 18.4% of the total fatty acids pool in cultivated samples, compared to <5% in native fat. This may result from residual bovine serum-derived lipids, donor-specific metabolic variability, or incomplete lipid remodeling during differentiation, suggesting further refinement of the culture medium could improve lipid specificity.

PUFA distribution (Figure 4C) also showed small but detectable levels of omega-3 (n-3) and omega-6 (n-6) fatty acids, likely originating from serum components. Notably, the presence of eicosapentaenoic acid (EPA, C20:5n3), a marine omega-3 fatty acid rarely found in terrestrial animal fat, is especially promising. Although present at low levels (0.3%), its inclusion suggests that cultivated fat could be tailored for health-targeted applications, such as omega-3-enriched meat analogues for individuals with limited fish consumption or increased clinical need [70].

Altogether, our fatty acid profile mirrors that of high-quality Wagyu beef. The high oleic acid content contributes positively to flavor release, tenderness, and nutritional appeal. The composition and structure of this engineered fat closely approximate native beef, including key MUFAs associated with sensory and nutritional quality. Moreover, comparisons with bADSCs cultivated in CMF/fibrin, our previous model [15], producing lower oleic acid (42%) and higher stearic acid (26%), highlight the influence of scaffold and mechanical environment on lipid biosynthesis. These findings establish a strong basis for further development of structured, functional, and health-enhancing cultivated fat for meat applications.

### Bioprinting possibility assessment of alginate bovine adipose tissue

To evaluate the applicability of the optimized alginate formulation for cultivated meat fabrication, its compatibility with our previously developed TIP (tendon-integrated printing) method [39] for structured meat assembly was next tested. Using gelatin-based support baths containing CaCl to induce alginate crosslinking, bADSC-laden fibers were successfully bioprinted (Figure 5A-D) and cultivated under adipogenic conditions. After 7 days, the bioprinted fibers demonstrated extensive maturation of bovine adipocytes, with a morphology and lipid content comparable to the ball tissues used previously in this study, as confirmed by Nile Red staining (Figure 5E,F). These results further validate the feasibility of using alginate-based bioinks for bioprinting high-fat cultivated meat structures.

**Figure 5.**
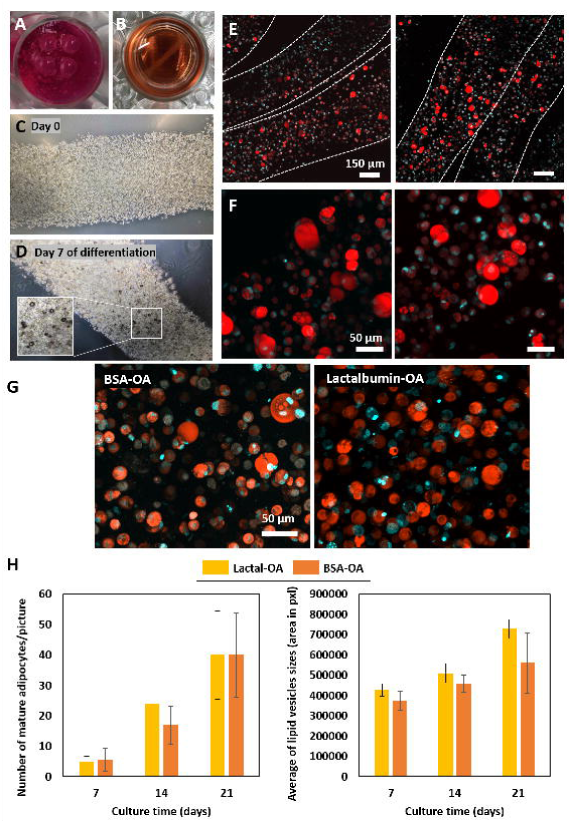
Bovine adipose tissue formation in 3D alginate gels and effect of lactalbumin–oleic acid formulations. (A–B) Top view of alginate-based fiber printed constructs just after printing (A) or after removing the gelatin supporting bath and cultured for 7 days differentiation (B) conditions in 6-well plates. (C–D) Phase-contrast representative images of bioprinted fibers at Day 0 (C) and Day 7 (D) of adipogenic differentiation. The inset in (D) highlights formation of lipid vesicles within the tissue. (E–F) Confocal representative microscopy images of bioprinted fibers adipose tissues cultured for 7 days under adipogenic conditions, stained with Nile Red and Hoechst. Dashed lines in (E) indicate printed fiber boundaries. (G) Confocal representative images of bovine adipocytes cultured for 21 days in 3D alginate under BSA–oleic acid (left) or lactalbumin–oleic acid (right) condition , stained with Nile Red and Hoechst. (H) Quantification of adipogenesis in BSA–OA and lactalbumin–OA conditions over 7, 14, and 21 days: left, number of mature adipocytes per image (vesicle diameter >35 µm); right, average lipid vesicle size (area in pixels). n = 3 biological replicates. Error bars represent mean ± SD.

### Towards a fully edible bovine adipogenesis reconstruction with further improvement by cell density and culture duration

To develop a fully edible albumin-based system for delivering oleic acid during adipogenic differentiation, we finally had to replace bovine serum albumin (BSA), which is not approved as a food additive in Japan, with edible alternatives such as α-lactalbumin or Whey Proteins (WP). Both α-lactalbumin alone and WP (including both α-lactalbumin and β-lactoglobulin) have known binding sites for fatty acids, α-lactalbumin exhibits one high affinity oleic acid binding site in its apo form, associated with other lower-affinity sites, and β-lactoglobulin contains hydrophobic cavities that accommodate long-chain fatty acids such as oleic and linoleic acids [71–73].

Using these proteins, we compared lipid accumulation in engineered bovine adipose tissues using as well longer differentiation duration culture. First, the use of lactalbumin-oleic acid condition instead of BSA-oleic was assessed by confocal imaging following Nile Red and Hoechst staining (Figure 5G, after 21 days of differentiation). It confirmed the intracellular lipid accumulation in both groups through the differentiation duration, displaying abundant, large unilocular lipid droplets with robust lipid storage, typical of mature adipocytes (Figure 5G). The mature adipocytes size was also compared (Figure 5H), revealing similar mature adipocytes numbers (lipid vesicle diameter >35 µm) content and whole similar size of lipid vesicles in the lactalbumin–oleic acid group compared to the BSA–oleic acid control. This suggests that lactalbumin is also capable of delivering oleic acid effectively to differentiating adipocytes.

Then, to further assess the suitability of dairy products-derived edible fatty acid carriers for oleic acid delivery during adipogenic differentiation, we compared the three culture conditions with BSA, lactalbumin and WP complexed with oleic acid over a 28-day differentiation culture period (Figure 6A). All three conditions supported progressive adipogenesis, as demonstrated by time-dependent lipid droplet formation visualized by Nile Red staining. Remarkably, both lactalbumin–oleic acid and WP–oleic acid conditions promoted lipid accumulation similar to that observed with the conventional BSA–oleic acid formulation, with particularly enhanced lipid storage visible by days 21 and 28.

**Figure 6.**
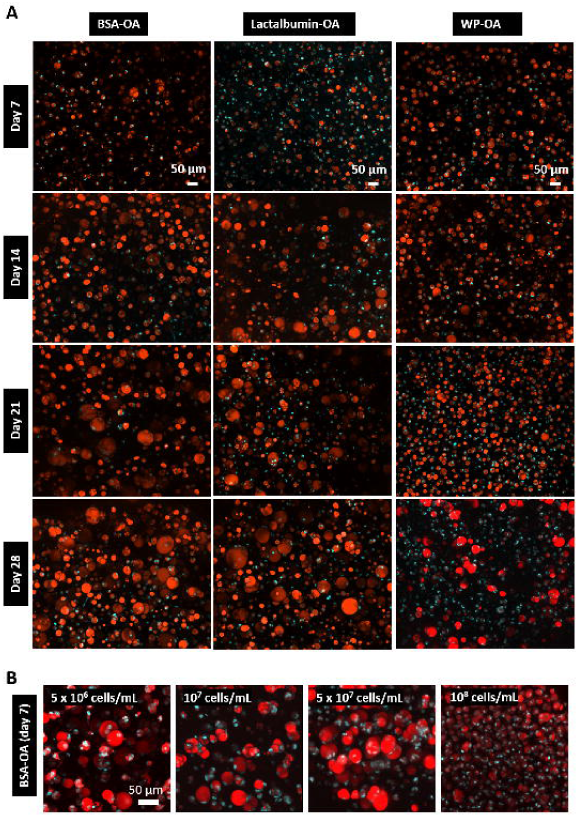
Time-dependent lipid accumulation and impact of cell density on adipogenesis using different protein-oleic acid complexes. (A) Confocal representative images of 3D bovine adipose tissues differentiated over 28 days alginate constructs with oleic acid complexed to BSA, lactalbumin, or whey protein. Nile Red and Hoechst staining were performed at Days 7, 14, 21, and 28. (B) Confocal representative images of 3D bovine adipose tissues with varying initial cell densities (5 × 10 , 1 × 10 , 5 × 10 , and 1 × 10 cells/mL) in alginate constructs and cultured for 7 days with BSA-oleic acid supplementation.

**Figure 7.**
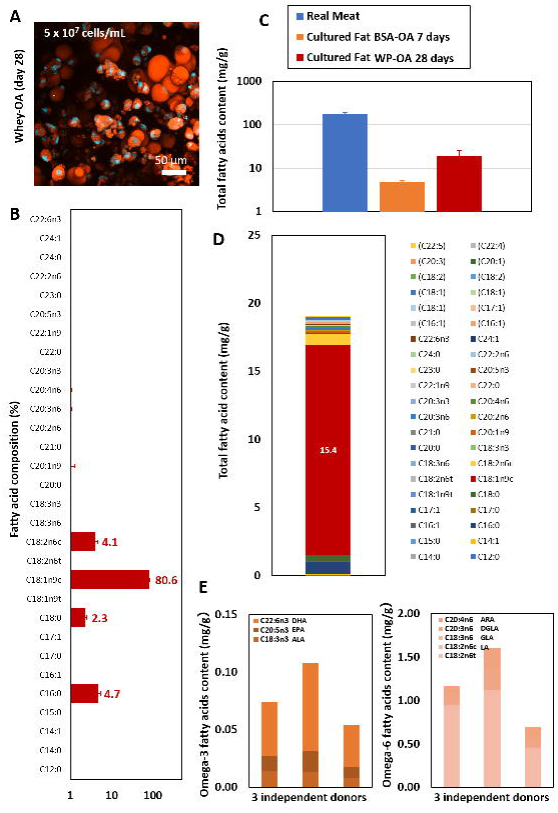
Fatty acid composition and quantitative lipid content of 3D cultured bovine adipose tissues differentiated for 28 days using whey-oleic acid supplementation. (A) Representative confocal image of lipid-rich 3D adipose alginate constructs stained with Nile Red and Hoechst after 28 days of differentiation culture with whey-oleic acid. (B) Relative fatty acid composition of the 28 days differentiation cultured fat. (C) Total fatty acid content (mg/g of total bovine fat tissue) in native bovine fat and cultured tissues. (D) Absolute content (mg/g of total bovine fat tissue) of the different fatty acids in the cultured fat samples. (E) Quantification of omega-3 (left panel) and omega-6 (right panel) fatty acid subtypes present in cultured samples, including ALA, EPA, DHA, LA, and GLA. N = 3 independent donors. Error bars represent mean ± SD.

Finally, after confirming the differentiation duration effect, we also investigated the effect of cell density to further enhance lipid accumulation in *in vitro* bovine adipose tissues (Figure 6B). In BSA-OA condition, embedding bovine ADSCs at a concentration of 5 × 10 cells/mL within alginate balls for 7 days of differentiation led to more pronounced adipogenic differentiation compared to both the lower density of 5 × 10 cells/mL used in earlier experiments and a higher density of 1 × 10 cells/mL, which exhibited diminished differentiation. One plausible explanation is that excessive cell packing at high densities may restrict intracellular lipid storage capacity, thereby impairing adipogenesis. Additionally, even though the culture medium volume was proportionally increased to match the cell number, the larger volume required at higher densities could exert additional mechanical forces on the tissues due to increased weight, potentially influencing cellular behavior and differentiation outcomes, especially adipocyte maturation. Sheng et al. [74] indeed reported that reduced medium volumes in adipocyte cultures promoted lipid accumulation and adipokine secretion, while larger volumes inhibited these processes. Physical parameters such as spatial constraints and mechanical loading may also play critical roles in regulating adipogenic efficiency and lipid accumulation [75–77].

Collectively, all these findings support the feasibility of replacing BSA with edible protein carriers for oleic acid delivery in adipogenic differentiation protocols and the use of 5 ×

## 10 cells/mL cell density for the further experiments

### WP–oleic acid supplementation enables reproduction of fatty acid profiles with enhanced similarity to native bovine fat

Compared to purified BSA or isolated lactalbumin, WP offer a more sustainable and cost-effective alternative, as they are readily available as by-products of cheese and milk processing, requiring minimal purification and thereby reducing overall environmental impact. Additionally, the broader spectrum of milk-derived proteins in WP may enhance the stability and bioavailability of oleic acid, supporting more efficient uptake and metabolic incorporation during adipogenesis. For these reasons, WP was selected for the final analysis of fatty acid composition and content.

To evaluate the lipid accumulation and fatty acid profile of bovine adipocytes cultivated in the alginate-based 3D system, we analyzed the cultivated fat tissues generated under WP-OA condition, after 28 days of culture at high cell density (5 × 10 cells/mL), where the adipocytes displayed mostly a multilocular morphology of high diameter size, as shown by Nile Red staining (Fig. 7A). The fatty acid composition of this 28 day differentiated cultivated fat was dominated by monounsaturated oleic acid (C18:1n9c), accounting for 80.6% of total fatty acids (Fig. 7B, individual results for each of the 3 independent donors in Supplementary Figure 2). While this proportion exceeds that typically found in native intramuscular fat (see Figure 4B), it further confirms the high degree of customizability of the *in vitro* culture system, demonstrating that adipocytes can be selectively driven to accumulate large amounts of oleic acid through targeted supplementation and culture conditions. This enrichment of oleic acid can be beneficial for further improving the rich umami flavor, tenderness, low melting point, and health profile of the final cultivated fat, especially when mixed with cultivated muscle content too. Palmitic acid (C16:0, 4.7%) and linoleic acid (C18:2n6c, 4.1%) can contribute to structural integrity and oxidative flavor brightness, respectively, while stearic acid (C18:0, 2.3%) can support mouthfeel and melting behavior. Minor fractions of myristic acid (C14:0, 0.4%) and palmitoleic acid (C16:1, 0.4%) might add subtle creaminess realism [78].

Quantitative analysis of the total fatty acid content (Fig. 7C) revealed that real meat samples still contained the highest lipid levels, as expected (around 180 mg/g tissue for real fat, compared with around 20 mg/g for cultivated fat at day 28, of average for 3 different donors). However, this long-term culture with WP-OA supplementation significantly enhanced lipid accumulation in the 3D cultivated fat, reaching 4-fold higher levels than those observed at day 7 with BSA-OA supplementation. This result highlights the importance of prolonged culture and optimized nutrient composition for effective adipogenesis and lipid maturation *in vitro*, even though real fat still contains 9-fold higher lipid content. Continued refinement of the culture conditions may gradually bridge this gap.

The absolute fatty acid content profile (Fig. 7D, individual results for each of the 3 independent donors in Supplementary Figure 2) shows again that oleic acid was the predominant fatty acid (15.4 mg/g), while other fatty acids such as palmitic (C16:0), stearic (C18:0), and linoleic acids were present in lower amounts. From this data, essential fatty acid subtypes were further analyzed separately, including omega-3 (Fig. 7E, left) and omega-6 fatty acids (Fig. 7E, right). Although the total omega-3 fatty acid content remained low (<0.15 mg/g), alpha-linolenic acid (ALA, C18:3n3) and trace levels of eicosapentaenoic acid (EPA, C20:5n-3) and DHA (C22:6n3) were detectable. On the other hand, omega-6 fatty acids were more abundant, particularly linoleic acid (LA, C18:2n6c, 0.8 mg/g on average), arachidonic acid (ARA, C20:4n6, 0.16 mg/g on average) and dihomo-γ-linolenic acid (DGLA, C20:3n6, 0.15 mg/g on average). This composition is consistent with a typical bovine adipose fatty acid profile. Although still present in only small amounts, the detection of EPA in the cultivated fat is a promising breakthrough, being rarely found in terrestrial animal products, yet it plays a critical role in reducing inflammation and promoting cardiovascular health [70]. Its presence highlights the transformative potential of cultivated meat not just to replicate conventional foods, but to reimagine them as vehicles for targeted health benefits. This opens the door to a new generation of functional, nutritionally enhanced meat alternatives, engineered to deliver specific bioactive compounds like EPA to populations with dietary restrictions or therapeutic needs, potentially positioning cultivated meat at the intersection of nutrition, sustainability, and preventive healthcare.

Interestingly, compared to native bovine fat, in general, the 28 days differentiated adipose tissue displayed a more diverse and functionally enriched fatty acid profile, especially for the long-chain polyunsaturated fatty acids (PUFAs) (these included linoleic acid (C18:2n6c), dihomo-γ-linolenic acid (C20:3n6), arachidonic acid (C20:4n6), and docosahexaenoic acid (C22:6n3)) (Supplementary Figure 3). This enrichment suggests that *in vitro* adipogenesis activates elongation and desaturation pathways [79,80] that are either suppressed or tightly regulated *in vivo*, possibly due to dietary constraints or ruminant-specific lipid metabolism. While this divergence may indicate incomplete terminal differentiation, it also introduces the potential nutritional advantages, as these PUFA fatty acids can be associated with anti-inflammatory, cardioprotective, and neuroprotective effects [81].

Comparing the distribution of saturated and monounsaturated fatty acids (from C12:0 to C18:0) can also reveals important insights into adipogenic progression and sensory properties across the conditions. Short and medium chain saturated fatty acids (C15:0, C16:0, C17:0 or C18:0) were found more abundant at 7 days of differentiation than at 28 days, suggesting an early burst of *de novo* lipogenesis that tapers as cells mature. This decline reflects the metabolic shift toward desaturation and elongation pathways or lipid remodeling during late-stage adipogenesis [38,62]. Notably, myristoleic acid (C14:1), a monounsaturated fatty acid derived from C14:0 via Δ9-desaturation, was absent at 7 days but emerged at 28 days, consistent with a probable upregulation of stearoyl-CoA desaturase (SCD1) activity arising during late-stage of adipocyte differentiation [82,83]. In contrast, heneicosanoic acid (C21:0) was predominantly found in real fat and nearly absent in both engineered conditions, highlighting a gap in very-long-chain fatty acid elongation under *in vitro* conditions.

These compositional dynamics might influence not only the biophysical properties of the lipid droplets, such as membrane fluidity and melting behavior, but also the sensory characteristics of the fat. Saturated fatty acids can contribute to firmness and mouthfeel, while monounsaturates like C14:1 and C16:1 can add creaminess and subtle flavor complexity [78,84]. Together, these shifts reflect a partially matured adipogenic state at 28 days, with potential for further optimization to balance nutritional value, textural realism, and flavor fidelity.

### Effect of FBS concentration on triglyceride storage in lactalbumin- and WP-supplemented adipogenesis

Finally, to investigate the possibility of reducing serum dependency in the adipogenic differentiation protocol, we examined the effect of lowering fetal bovine serum (FBS) concentrations on triglyceride accumulation in bovine ADSC-derived adipocytes cultivated with either lactalbumin or WP supplementation (Figure 8). In this experiment, FBS was used instead of edible bovine serum solely to evaluate the effect of serum reduction, as our previous study demonstrated no significant difference between DMEM + FBS and IMEM + edible bovine serum for both cell proliferation and differentiation efficiency [15]. Therefore, these results can be extrapolated to edible serum–based conditions.

**Figure 8.**
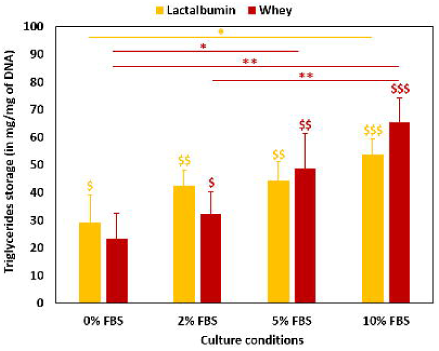
Triglyceride accumulation in bADSC-derived adipose tissues cultured with whey- or lactalbumin-oleic acid (OA) complexes under varying FBS concentrations. Quantification of intracellular triglyceride content normalized to DNA after 7 days of adipogenic differentiation in the presence of 0%, 1%, 5% or 10% FBS. Comparison between whey-oleic acid and lactalbumin-oleic acid supplementation conditions. Error bars represent mean ± SD; N = 3 independent donors. $ means statistical comparison with undifferentiated condition. Asterisk (*) denotes significant statistical enrichments (\**p*<0.05, ** *p*<0.01 and *** *p*<0.001).

First, the amount of triglycerides detected after 28 days of differentiation in the WP-OA culture condition, normalized to DNA content (Figure 8), was 86-fold higher than that measured previously in the OA-only condition after 7 days of differentiation (Figure 2B). This substantial increase highlights the enhanced lipid accumulation achieved by switching to albumin-complexed OA, increasing cell density, and extending the differentiation period. These findings further confirmed that albumin-complexed OA improved fatty acid delivery and cellular uptake and metabolic processing, to promote adipogenic maturation. Then, in both conditions, lipid storage increased progressively with higher FBS concentrations, indicating that serum indeed plays an essential role in adipogenesis in this system. In lactalbumin-supplemented cultures, TG accumulation rose with increasing FBS but showed no significant difference between 2% and 10% FBS, suggesting that near-maximal lipid storage can be achieved with reduced serum levels. In contrast, WP supplementation supported a more gradual TG accumulation, with significant differences observed between 2% and 10% FBS. Under these conditions, lipid storage at 5% FBS remained intermediate, indicating that further reduction below this concentration can compromise adipogenesis. The WP complex protein mixture does not fully replace the role of serum components. Nevertheless, from an application perspective, reducing FBS or edible bovine serum from the differentiation protocol could reduce cost, ethical concerns, and batch variability, making the WP-OA formulation highly attractive for future research on scalable and FBS-free cultivated fat production.

## Conclusion

This study presents a robust and fully edible strategy for bovine adipogenesis that advances the development of functional cultivated fat tissues for food applications. We demonstrated that replacing animal-derived scaffolds such as fibrin with algae-derived alginate enables stronger adipogenic differentiation while maintaining physiologically relevant mechanical properties. Among the edible protein carriers evaluated, WP protein emerged as a suitable edible alternative to both BSA and lactalbumin for oleic acid delivery, promoting efficient lipid accumulation even under lower serum content conditions. This finding is particularly significant for the cultivated meat field, where reducing bovine serum is a major step toward regulatory compliance, ethical acceptability, and cost reduction.

WP, which are by-products of the dairy industry, offer a sustainable, food-grade, and nutritionally rich solution that supports both adipogenic commitment and terminal adipocyte maturation, notably with an accumulation of high levels of monounsaturated fatty acids, especially oleic acid, which are key to flavor, texture, and nutritional value in premium beef such as Wagyu. These results underscore WP’s dual role as both a functional nutrient carrier and a bioactive culture medium component, aligning with recent advances in food-grade cell culture systems and circular bioeconomy models.

Furthermore, our engineered adipose tissues demonstrated rheological properties, fatty acid profiles, and lipid droplet morphologies that closely resemble native bovine fat. While total lipid content remains lower than *in vivo* tissues, long-term culture and optimized cell densities substantially improved adipogenic outcomes, pointing to the tunability of *in vitro* fat constructs for desired sensory and nutritional characteristics, tailored for food innovation. Future optimization can focus on metabolic modulation, mechanical stimulation, and dynamic culture systems such as perfusion bioreactors to further enhance lipid yield and tissue maturation.

Beyond meat alternatives, the oleic acid–enriched adipose tissues developed in this study could serve as customizable platforms for functional food applications, including the delivery of health-promoting fatty acids such as omega-3s. By leveraging edible biomaterials and dairy-derived protein carriers, this approach advances the vision of next-generation cultivated foods that are not only ethically and environmentally sustainable, but also nutritionally enhanced and organoleptically satisfying.

