## Supplementary Figures for "Meat the Fat: From High Lipid Content to Nutritional Enhancement to Reconstruct Wagyu Fat with Tunable Rheology and Organoleptic Fidelity for Cultivated Meat"

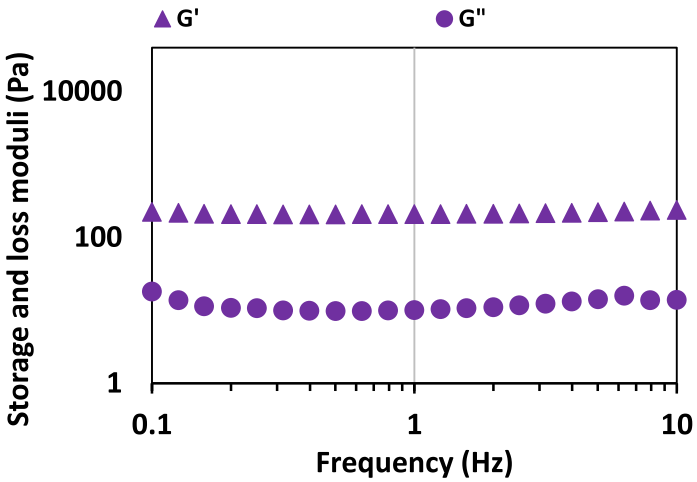


**Supplementary Figure 1.**
Storage modulus (G′) and loss modulus (G″) of cell-free 1% alginate gels assessed by frequency sweep rheometry after 9 days of culture with medium change every 2-3 days.

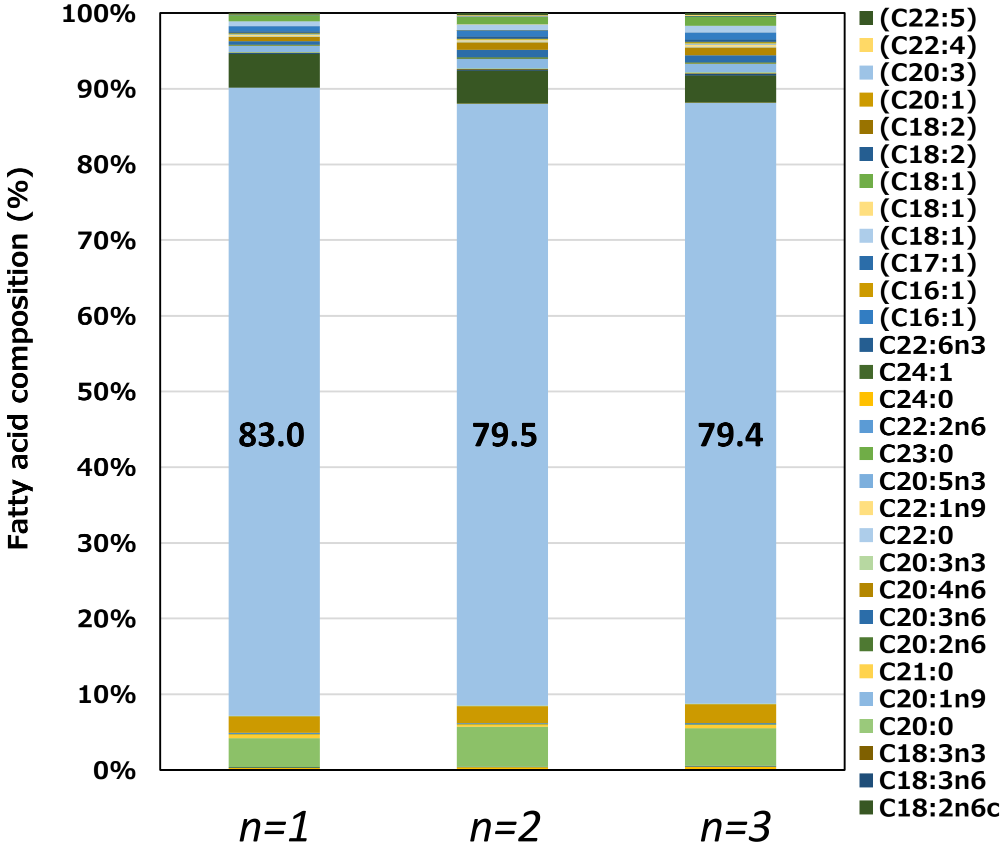


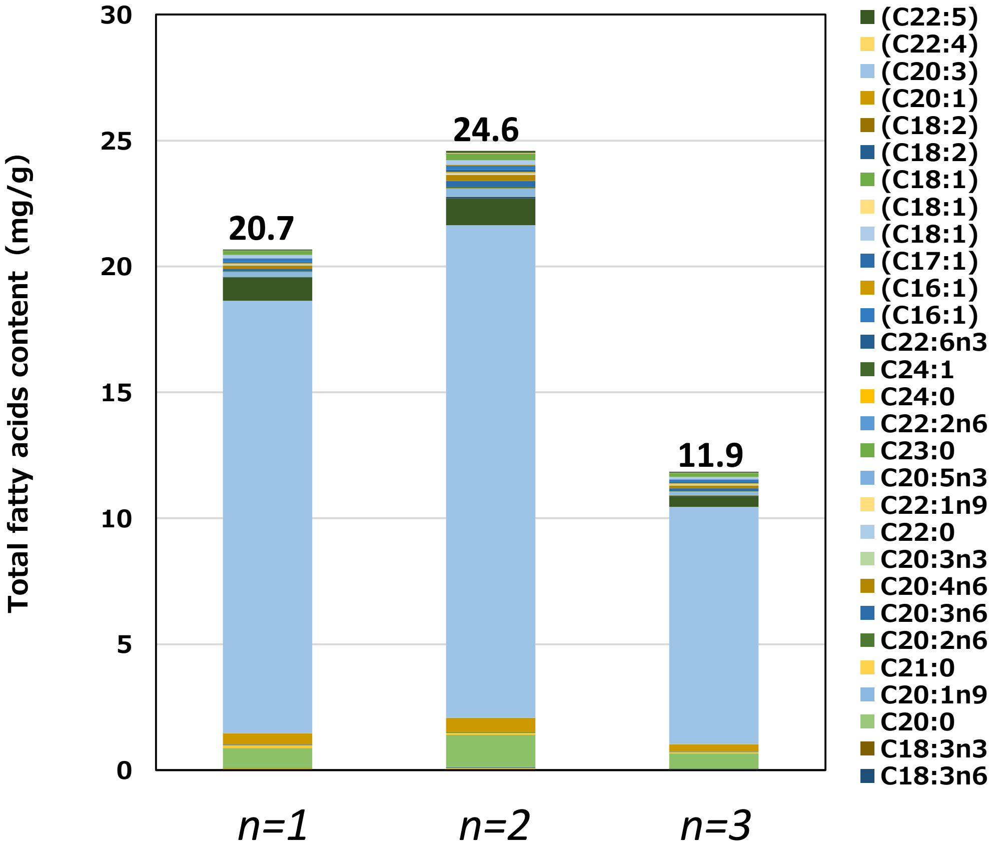


**Supplementary Figure 2.**

(top) Relative fatty acid composition of the 28 days differentiation cultured fat with Whey-OA (3 independent donors).

(bottom) Total fatty acid content (mg/g of tissue) of the 28 days differentiation cultured fat with Whey-OA (3 independent donors).

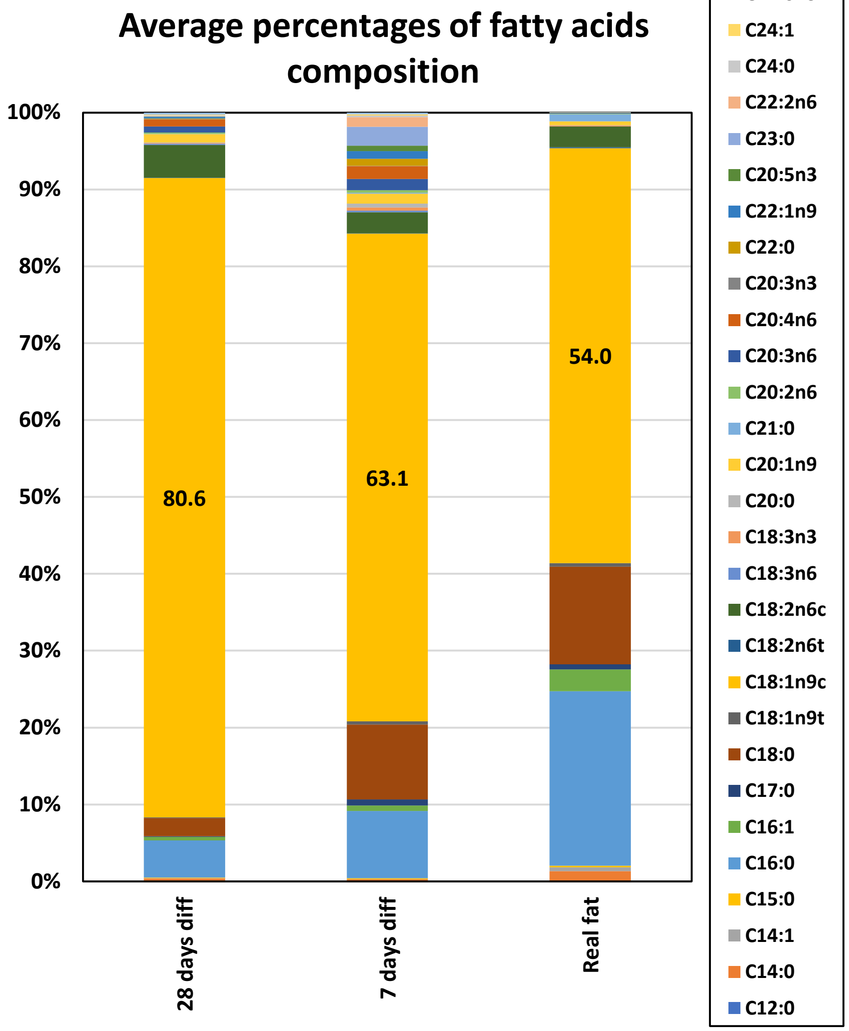


**Supplementary Figure 3.** (top) Relative fatty acid composition in percentages of the 28 days differentiation cultured fat with Whey-OA, the 7 days differentiation cultured fat with BSA-OA and the real bovine fat comparison (averages from 3-6 independent donors). (bottom) Repartition graphical representation of each fatty acid content for each condition.

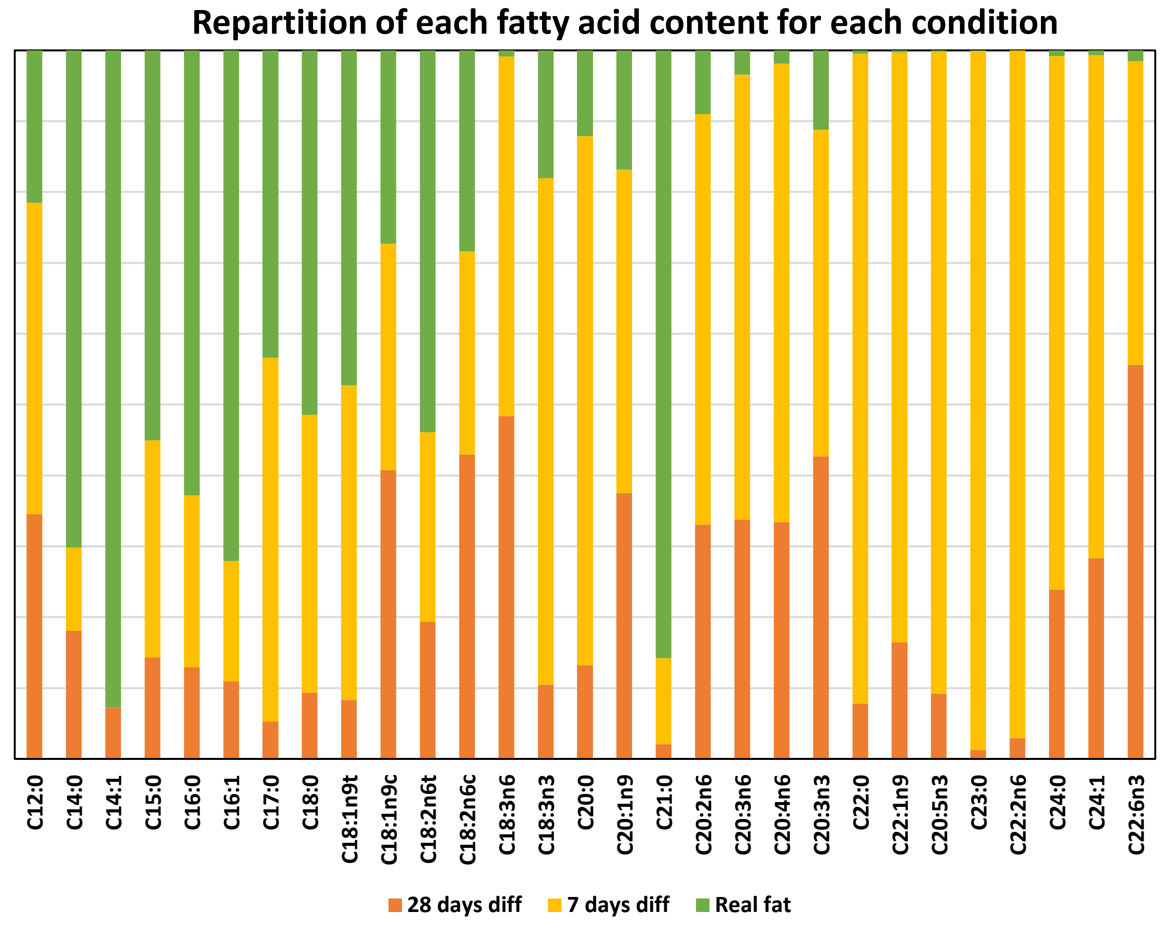
